# Temporal and spatial changes in phyllosphere microbiome of acacia trees growing in super arid environments

**DOI:** 10.1101/2020.01.02.893446

**Authors:** Ashraf Al-Ashhab, Shiri Meshner, Rivka Alexander-Shani, Michael Brandwein, Yael Bar Lavan, Gidon Winters

**Author notes:** Corresponding author: Dr. Ashraf Al Ashhab.

## Abstract

Along the arid Arava, southern Israel, acacia trees (*Acacia raddiana* and *Acacia tortilis*) are considered keystone species. In this study, we investigated the ecological effects of plant species, microclimate (different areas within the tree canopies) and seasonality on the endophytic and epiphytic microbiome associated with these two tree species. 186 leaf samples were collected along different seasons throughout the year and their microbial communities were studied using the diversity of the 16S rDNA gene sequenced on the 150-PE Illumina sequencing platform. Results show that endophytic, but not epiphytic, microbiome communities were different between the two acacia species. Endophytic, but not epiphytic, microbiome was affected by temporal changes (seasons) in air temperature. Acacia canopy microclimate was also found to have a significant effect on exosphere microbiome, with *A. tortilis* having a higher microbial diversity than *A. raddiana* with significantly different community compositions in different seasons.

**Importance:** The evolutionary relationships and interactions between plants and their microbiome are of high importance to the survival of plants in extreme conditions. Changes in microbiome of plants can affect plant development, growth and health. In this study, we explored the relationship between keystone desert trees and their microbiome along seasonal variation. These results shed light on the importance and uniqueness of desert phyllosphere microbiome. Although acacia trees are considered keystone species in many arid regions, to the best of our knowledge, this is the first time that microbial descriptors have been applied in these systems. This work constitutes a new approach to the assessment of these important trees and a stepping stone in the application of microbial communities as a putative marker in a changing environment.

## 1. Introduction

The above-ground surfaces of plants (the phyllosphere) harbor a diverse variety of microorganisms, including bacteria. The plant phyllosphere microbiome has been shown to play an important role in the adaptation of the plant host to different environments such as tolerance to heat, cold, drought and salinity (1–5). Many researchers have demonstrated different desert plants eco-physiological adaptation to play a key role in microbial functional diversity (2, 6). While the exact correlation of phyllosphere microbial communities and these unique adaptations ae yet to be clarified, growing findings indicating that each plant type provides a suitable and unique microenvironment. Recent study investigated the adaptation of three Negev desert plant species finding Bacteroidetes to dominated the leaves of *H. scoparia* for instance and were not abundant in the other species they investigated (7). Plants phyllosphere microbes were also found to differ among different habitat and climate conditions comparing between arid, semi-arid and temperate habitat (8). These microbes were also found to correlate with high temperature, droughts and UN radiation (9, 10), regardless of their geographical distance (11). In this context, Desert phyllosphere microbiome showed to intervene in plant growth and alternative ways in the metabolism of some nutrients such as: fixing nitrogen (N) from atmospheric sources (1, 12), or by utilizing phosphorus (P) through solubilizing enzymes (13, 14) and by production of Siderophores molecules to bind iron (15, 16). Other phyllosphere microbiome also found to provide a defense against other bacteria pathogens such as blight disease (17), botrytis fungal infection (18).

Alongside seasonality (6, 19, 20) and canopy structure (21), plants showed to shape the phyllosphere microbiome. These studies indicated that both abiotic (climate-related) and biotic (plant genotype) factors play a pivotal role in structuring the phyllosphere microbial communities (22). Infact, endophytic (“inside the leaves of plants”) and epiphytic (“outside the leaves of plants”) microbial communities showed too different in the microbial community composition, were epiphytic bacterial communities were richer and more abundant compared to the endophytic bacterial communities, moreover, abiotic factors shown to have different effect on endophytic and epiphytic bacterial communities. Season was the major driver of community composition of epiphytes while wind speed, rainfall, and temperature were the major drivers for endophytic composition (23).

These complex interaction between plant microbiome (both endophytic and epiphytic) and different biotic and abiotic conditions within arid ecosystems, is of particular interest considering the current scenarios of climate change and desertification (24). Additionally, studies on microbiomes in arid plants could shed new light on important key microbial groups that might be of potential use in arid agricultural practices, biotechnology and plant adaptation strategies to climate change (25). In this study, we focused on the Negev desert (Fig. 1A) investigated the endophytic and epiphytic microbiome associated with the phyllosphere (leaves) *Acacia raddiana Savi* and *Acacia tortilis* (*Forssk*.) *Hayne* (Synonyms, respectively of *Vachellia tortilis* subsp. *raddiana* (*Savi*) and *Vachellia tortilis (Forssk.)*; (25; Fig. 1B).

**Figure 1.**
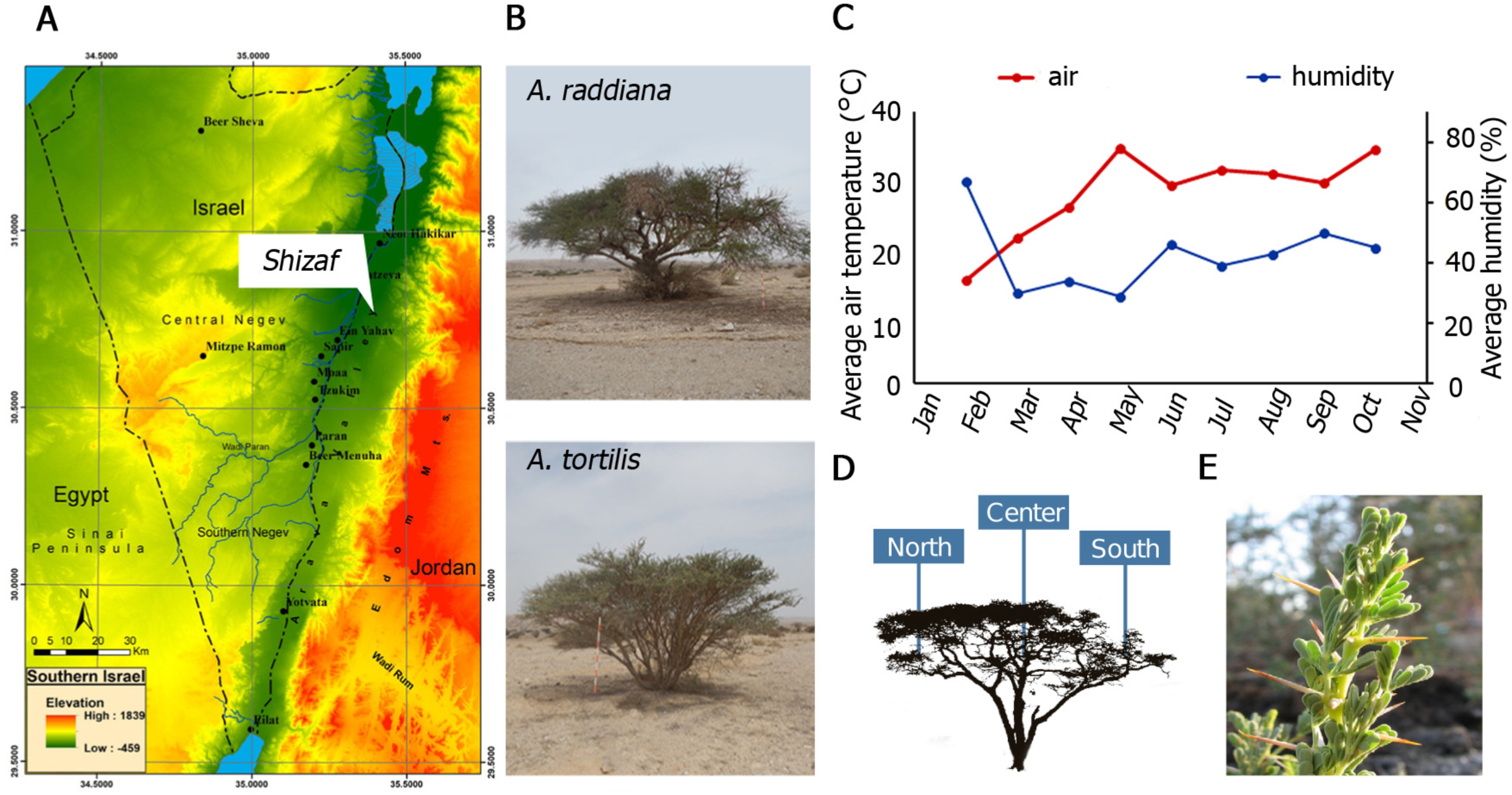
South Israel topography map showing the studied site of Wadi Sheizaf (A), and acacia trees (*A. raddiana* and *A. tortilis*; B) sampled monthly during 2015. Air temperature and humidity were hourly obtained from the Hazeva meteorological station (C). In each month, leaf samples (example of leaves collected shown in E) were collected from the north, center and south sides of the canopies (D). with a close-up picture showing the collected leaves for endophytic and epiphytic microbial community analysis.

These two tree species are found growing in some of the hottest and driest places on Earth. Within the arid Arava valley along the Syrian-African transform (Great Rift valley) in southern Israel and Jordan, *Acacia raddiana* and *Acacia tortilis* are the two most abundant and, in many places, the only tree species present (27). In these arid habitats, acacias are found mostly growing in the channels of ephemeral river beds (“wadis”, a term from Arabic; (28)). Both *Acacia raddiana* and *A. tortilis* are considered keystone species that support the majority of the biodiversity surrounding them and locally improve soil conditions for other plant species (28–32). We hypothesized that variations in the bacterial communities of phyllosphere would be associated not only with the host species (*A. raddiana* and *A. tortilis*), but also with sampling mporal changes), tree microclimate (leaves growing on the north or south side of tree at are exposed to direct sun radiation vs. shaded leaves).

## 2. Results

A total of 186 acacia leaves sample were collected for both epiphytic and endophytic microbial communities. After data processing, each of the five primer sets were examined separately for their coverage across the samples and their retained sequence numbers (Table S4).

Results show that when using the third primer set, we were able to retain the largest number of sequences for all the samples 13,944±13,710 (n=186, min=122) compared to the rest of the primers (Table 1). Thus, we based all our further analysis of bacterial communities on this primer set.

**Table 1.** Sequence number of the split files based on the different primer set with some basic statistics including sequences average ±SD, number of final samples after curation (n), and the minimum sequence number in each sample (min).

|  |  |  |
| --- | --- | --- |
| First primer set | 15,812±14,109 (n=186) | 4,569±6,594 (n=175, min=102) |
| Second primer set | 1,540±3,631 (n=186) | 589±961 (n=75, min=104) |
| Third primer set | 36,451±18,831 (n=186) | 13,944±13,710 (n=186, min=122) |
| Forth primer set | 7,774±8573 (n=186) | 1,565±3,513 (n=145, min=100) |
| Fifth primer set | 998±2,389 (n=183) | 1,441±2,434 (n=40, min=104) |

### 2.1. Acacia bacterial community composition of endophytic compared to epiphytic

The diversity estimates of epiphytic and endophytic bacterial communities, for both *A. raddiana* and *A. tortilis* at South “S” canopy sides are shown in Table 2. For all diversity estimates, the diversity of endophytic bacteria diversity was half of that found on the exosphere (Table 2) indicating a different bacterial community and diversity pattern that exists in these two microbial communities.

**Table 2.** Average diversity estimates (±SD) measured across the entire sampling campaign for south side (S) of the tree canopy for the epiphyte and endophyte microbial communities of *A. raddiana* and *A. tortilis*, observed number of OTUs, Chao1 species’ diversity estimate, microbial communities’ phylogenetic diversity and the Shannon bacterial communities’ diversity. Diversity metrics were calculated in QIIME-1 software.

| Species | Canopy | Observed<br>number of<br>OTUs | Chao1<br>species'<br>diversity<br>estimate | microbial<br>communities<br>phylogenetic<br>diversity | Shannon<br>bacterial<br>communities'<br>diversity |
| --- | --- | --- | --- | --- | --- |
| *A. raddiana* | *S-epiphyte* | 375±158 | 644±284 | 18.5±5.7 | 5.2±1.1 |
|  | S-endophyte | 171±41 | 350.3±70 | 10.3±2.1 | 2.7±0.5 |
| *A. tortilis* | *S-epiphyte* | 410±164 | 702±277 | 20.4±6.5 | 5.0±1.1 |
|  | S-endophyte | 135±35 | 275.0±73 | 8.7±2.1 | 2.7±0.7 |

To compare the diversities of epiphytic and endophytic bacterial communities extracted from leaf samples, acacia samples from south canopy sides (Table 2) were analyzed and plotted using NMDS based on Bray-Curtis distance matrix (Fig. 2). Two separate clusters for endophytic and epiphytic bacterial communities (Fig. 2A) were found to be significantly different (p=0.005). However, while both acacia species (*A. raddiana* and *A. tortilis*) demonstrated separate clusters within the endophytic bacterial communities (p-value=0.006, Fig. 2A and B), they did not separate into different clusters in the epiphytic samples (p-value=0.585, Fig. 2A). To illustrate these differences, major bacterial phyla were plotted for both species in exo- and endophytic samples (Fig. 3). Exosphere samples showed significantly higher abundance of Actinobacteria, Cyanobacteria and significantly lower abundance of Firmicutes and Proteobacteria compared with endophytic samples from the same leaves. While exosphere bacterial communities showed no significant changes in phylum composition between the host species (*A. raddiana* or *A. tortilis*), the endophytic bacterial communities differed between acacia species (Fig. 3). In endophytic bacterial communities, abundance of Firmicutes was significantly higher on *A. raddiana* compared with leaves sampled from *A. tortilis* trees (61.2 ± 32.0% and 32.0 ± 27.9%, respectively), while *A. tortilis* had a significantly higher abundance of Proteobacteria than *A. raddiana* (60.9 ± 26.4% and 27.7 ± 21.3%, respectively). Most Proteobacteria (>88%) belonged to *Comamonadaceae* family and most Actinobacteria (>90%) belonged to *Bacillaceae* family, while all had an unknown subfamily classification. Nevertheless, both *Bacillaceae* and *Comamonadaceae* families were dominated by one OTU at 97% cluster similarity, indicating similar bacterial OTUs that inhabit both *A. tortilis* and *A. raddiana* but at different proportions.

**Figure 2.**
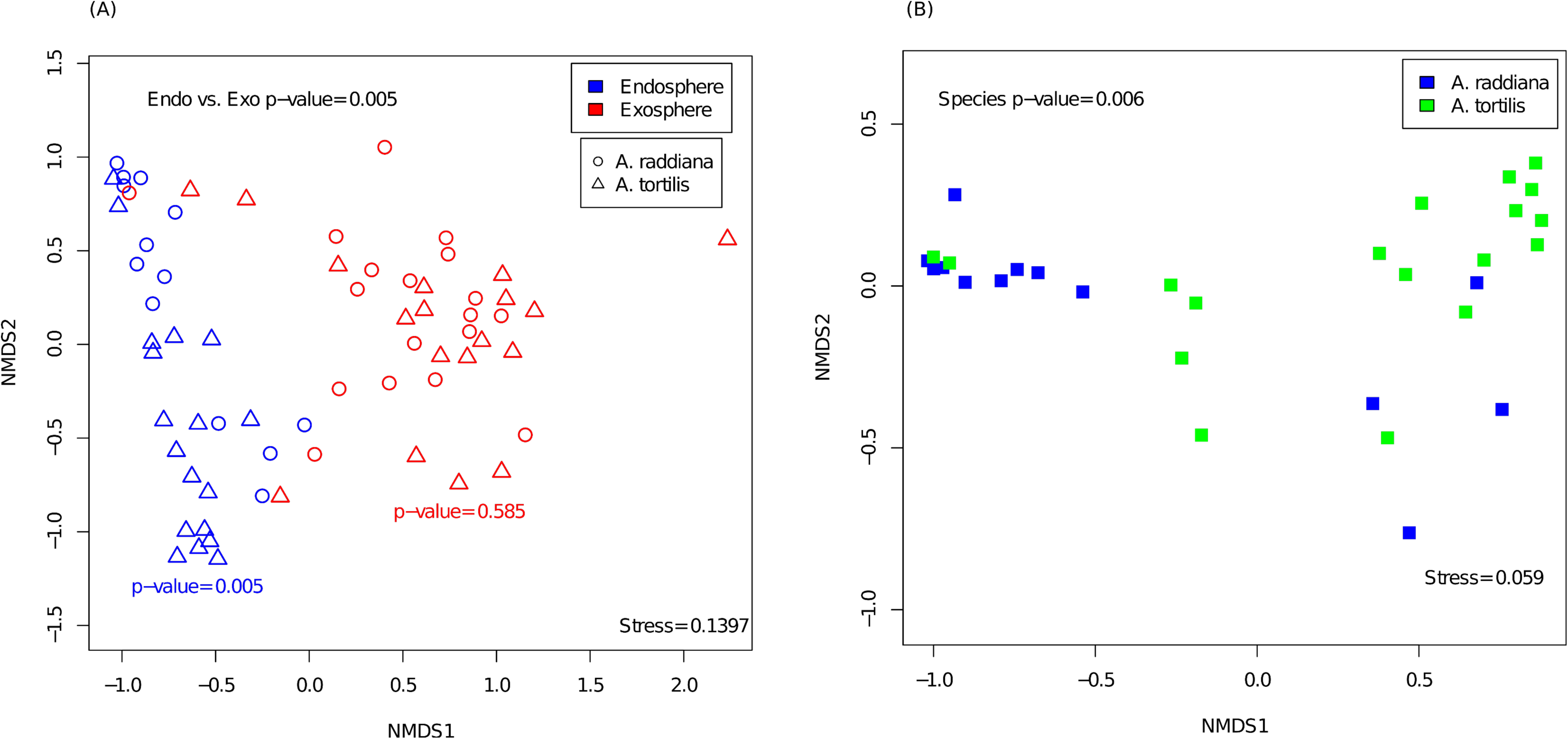
NMDS illustrating the phyllosphere bacterial community showing separate clusters of bacterial communities between (A) the epiphytic (red) and the endophytic (blue) bacterial communities from leaves sampled from south side canopy areas and (B) different clusters for *A. raddiana* (blue) and *A. tortilis* (green) for endophytic bacterial communities.

**Figure 3.**
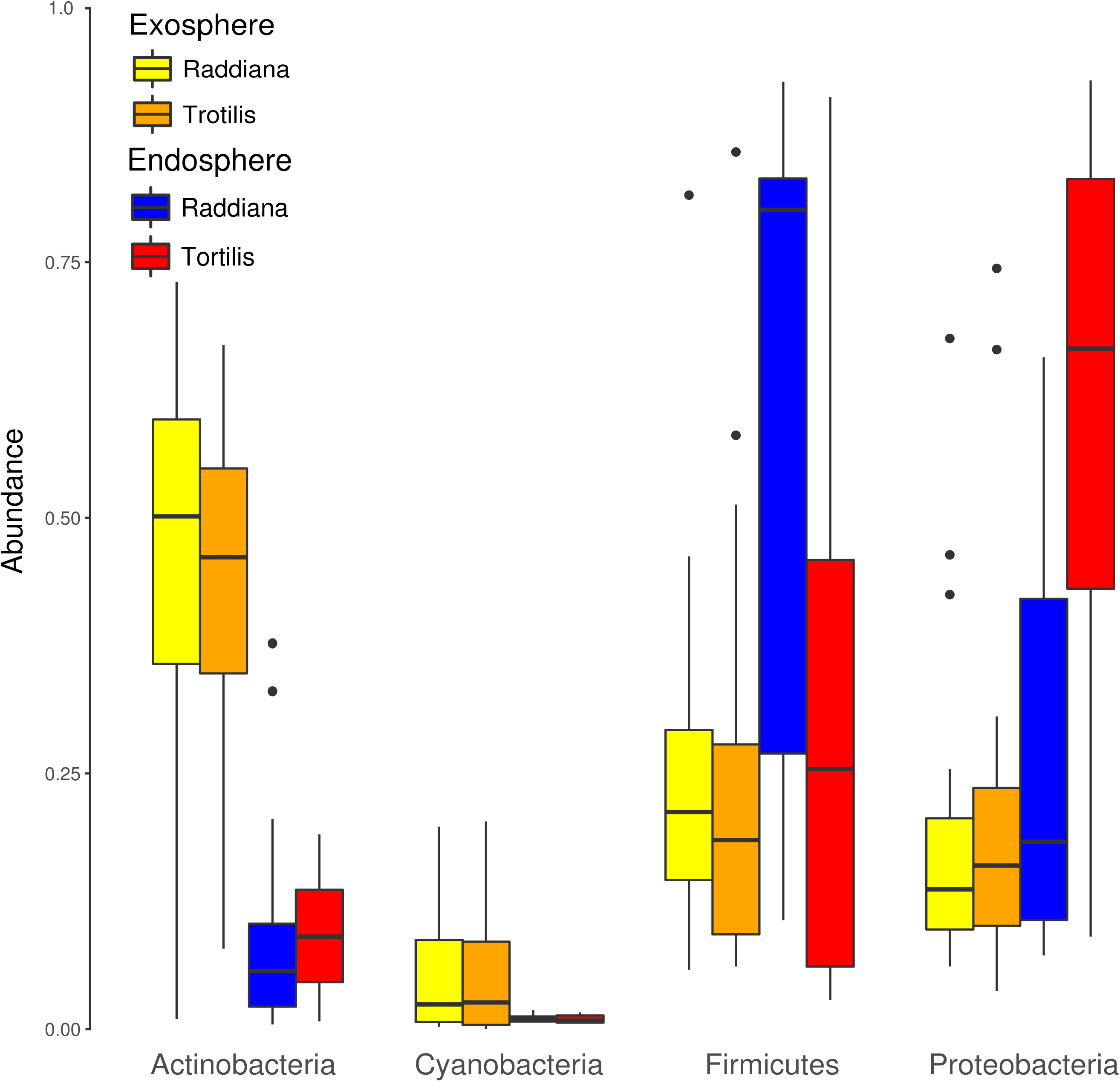
Box plot illustrating exo- and endosphere major bacterial phyla.

### 2.2. Acacia temporal and canopy variation of phyllosphere bacterial communities

To check the temporal effect on exo- and endophytic bacterial communities, acacia leaf samples from different sampling campaigns (Months; Table S1) were analyzed and plotted using NMDS based on Bray-Curtis distance matrix. Results (Fig. 4A) showed separate clusters for exosphere bacterial communities at different sampling months (p-value=0.001). For endophytic bacterial communities (Fig. 4B), different sampling months showed no clear separation (p-value=0.574), nevertheless a separate cluster was noticed for the samples collected in July-September and November compared with the endophytic bacterial communities collected in January, February, and in April-June (Fig. 4B).

**Figure 4.**
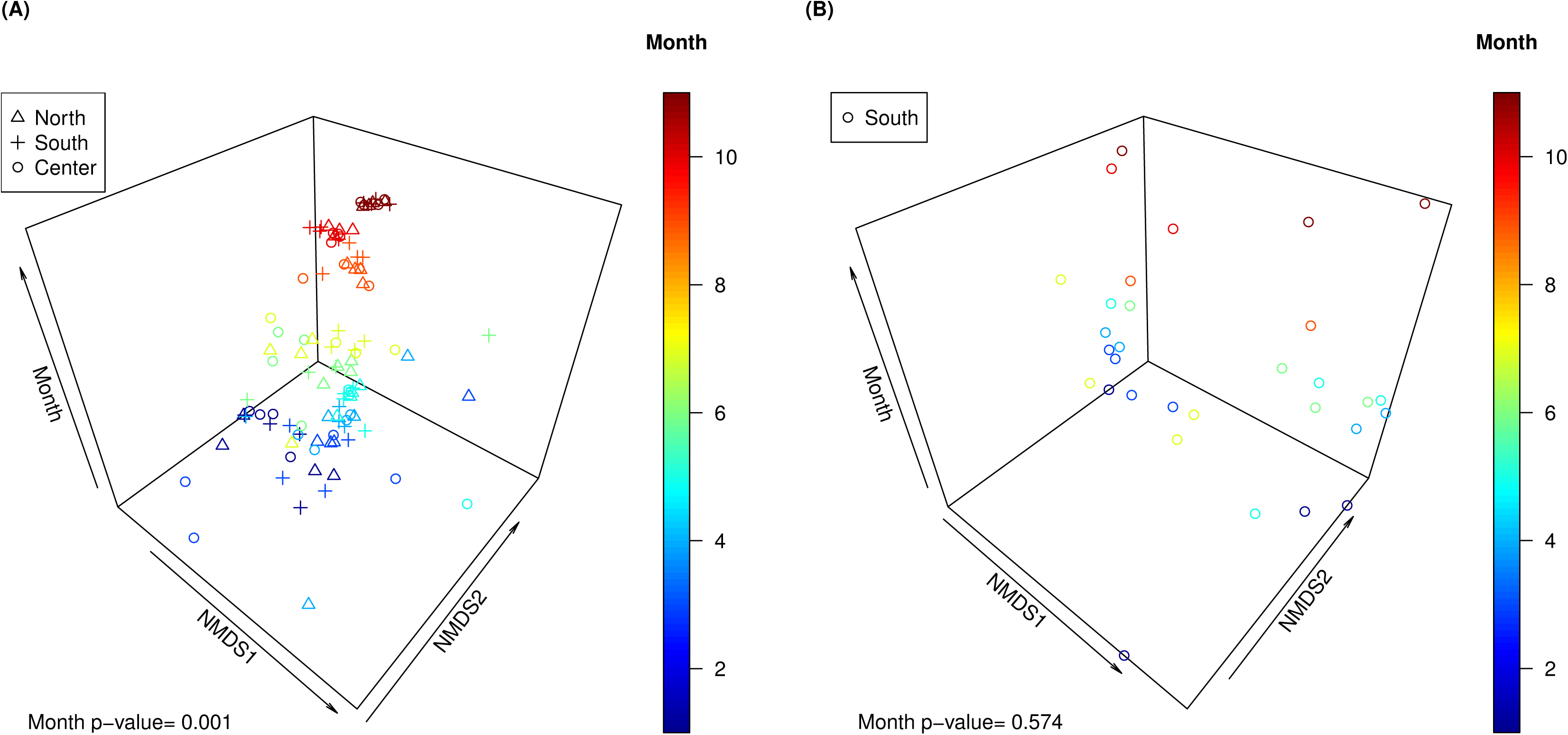
NMDS illustrating the phyllosphere bacterial communities showing separate clusters of bacterial communities between different sampling campaigns (Months) for (A) epiphytic and (B) endophytic bacterial communities.

To investigate the effect of microclimate (different canopy sides of the tree) on the epiphytes bacterial communities, the diversity estimates of epiphytic bacterial communities for both *A. raddiana* and *A. tortilis* in (i) north canopy side, (ii) center canopy shaded area were, and (iii) south canopy side compared (Table 3). Diversity estimates showed no clear differences between the different canopy sides for both *A. tortilis* and *A. raddiana*.

**Table 3.**
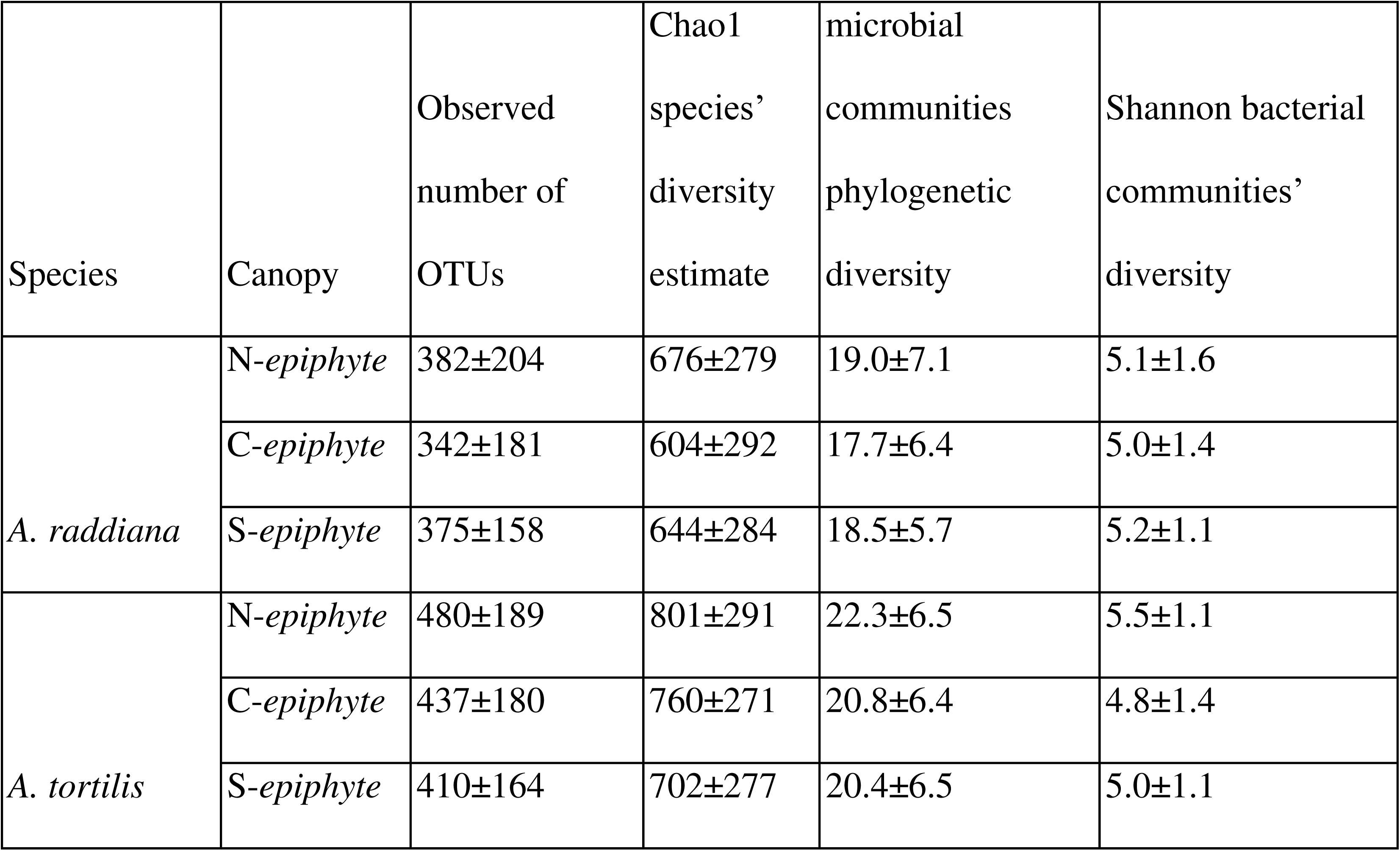
Average diversity estimates ± SD across the entire sampling campaign for epiphytic bacterial communities of *A. raddiana* and *A. tortilis*. Shown are the different canopies from which samples were taken from (N=north canopy side, C = center canopy shaded area, S= south canopy side), observed number of OTUs, Chao1 species’ diversity estimate, microbial communities’ phylogenetic diversity and the Shannon bacterial communities’ diversity. Diversity metrics were calculated in QIIME-1 software.

To understand what shapes the epiphytes bacterial diversity in different canopy sides, an NMDS cluster analysis was generated (Fig. 5). Result illustrated the bacterial community composition of the different canopy side (north, center or south) for *A. raddiana* and *A. tortilis*, showing no significant differences in the epiphytes bacterial communities between the different canopy sides (p-value=0.728), nor between the two species (*A. raddiana* and *A. tortilis*) (p-value=0.123), indicating a similar epiphytic bacterial diversity across the different sides of the trees.

**Figure 5.**
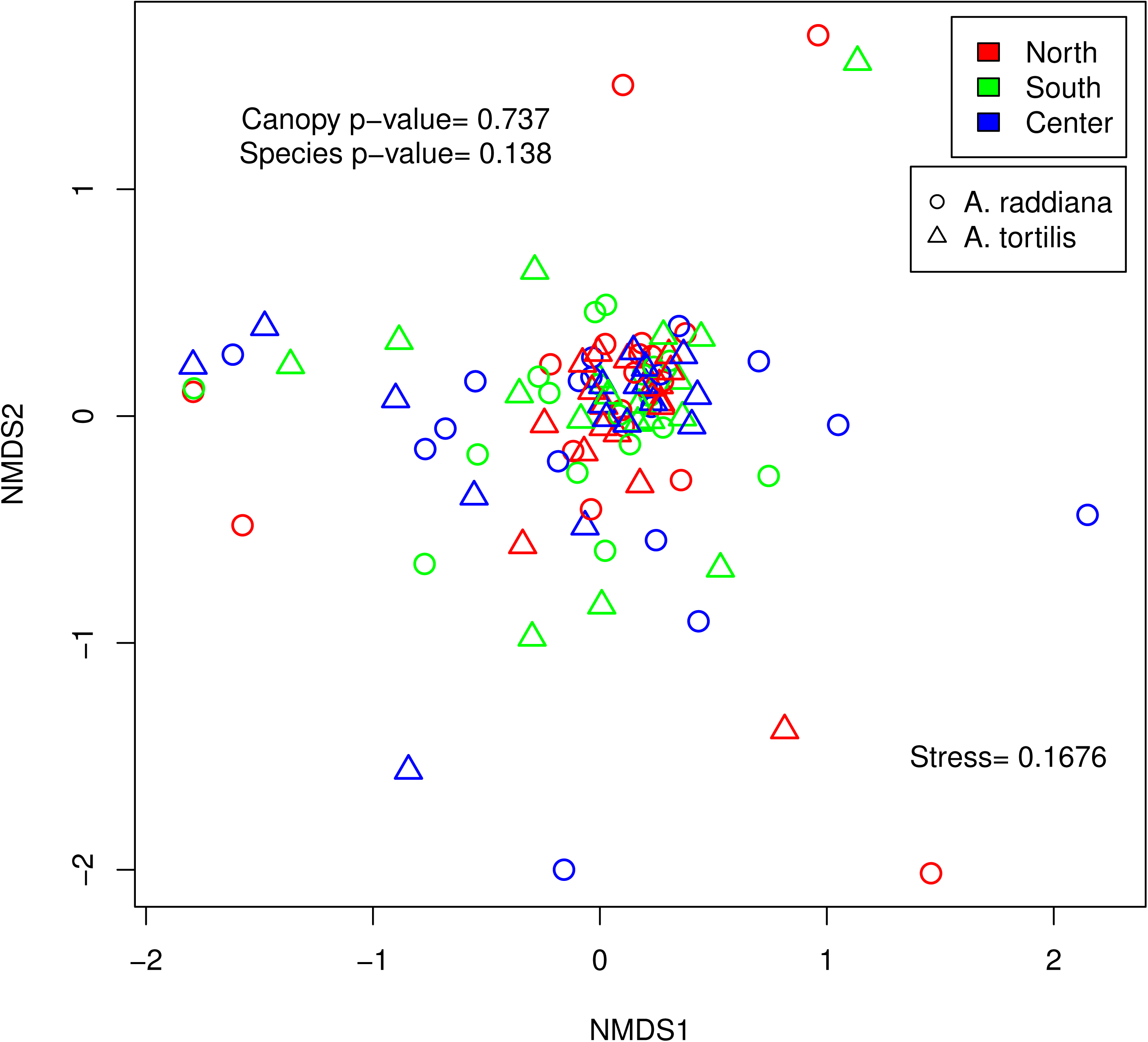
NMDS illustrating the epiphytes bacterial community at north (red), south (green) and center (blue) canopy side, for *A. raddiana* (circles) and *A. tortilis* (triangles).

To better illustrate different epiphytes bacterial phyla and their seasonality changes, we plotted the different bacterial phyla composition along with the different sampling months and canopy sides (Fig. 6). Temporal fluctuations were found between Actinobacteria and Firmicutes compositions, with the phyla Firmicutes found to be mostly dominant in January and July for both Center and South canopy sides (72.4 ± 14.6%), but not in the North canopy side which was dominated by Actinobacteria phyla (51.2 ± 17.7%) (Fig. 6A). Different patterns of bacterial diversity were also found in July, where both North and South were dominated by Actinobacteria (52.7 ± 5.8%) compared to the center canopy side that was dominated by Proteobacteria (70.2 ± 21.7%). In September, different clusters formed in the South, North and Center canopy sides, and all these canopy sides were dominated mostly by Acidobacteria (57.0 ± 8.7%) and Firmicutes (23.5 ± 5.5%). It should also be noted that in September, the different canopy sides hosted more similar proportions of these dominating bacterial groups.

**Figure 6.**
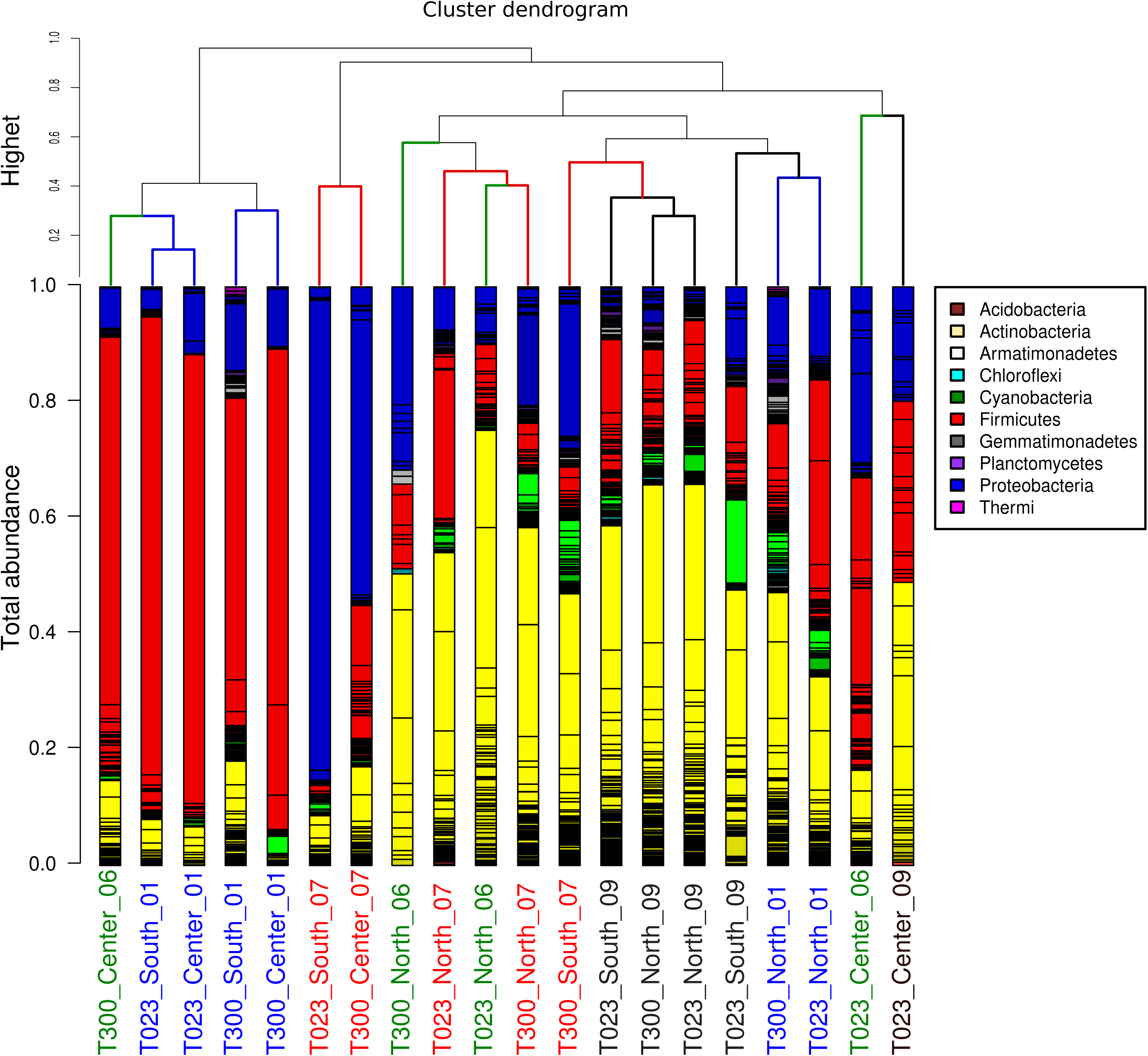
Epiphytes bacterial community composition clusters arranged primarily by sampling month and canopy side. A) Clustering dendrogram of bacterial communities, and (B) bacterial community composition for the samples shown in panel A. Different bar colors represent different phylum while shades of the same color represent different OTU’s. The X-axis titles were color coded for different month, January in blue, June in green, July in red and September in black.

To test whether other abiotic factors effect on the microbial communities differently for canopy sides, canonical correspondence analysis (CCA) (33) was performed for the epiphytic (Fig. 7A) and endophytic (Fig. 7B) bacterial communities of *A. raddiana* and *A. tortilis*. Only those abiotic factors with significant values (p-value ≦ 0.005) were plotted. Results show that temperature, 0.005) were plotted. Results show that temperature, VPD, humidity and precipitation had a significant effect on the epiphytes bacterial communities regardless of the canopy side (Fig. 7A), while temperature and precipitation had a significant effect on the endophytic bacterial communities (Fig. 7B).

**Figure 7.**
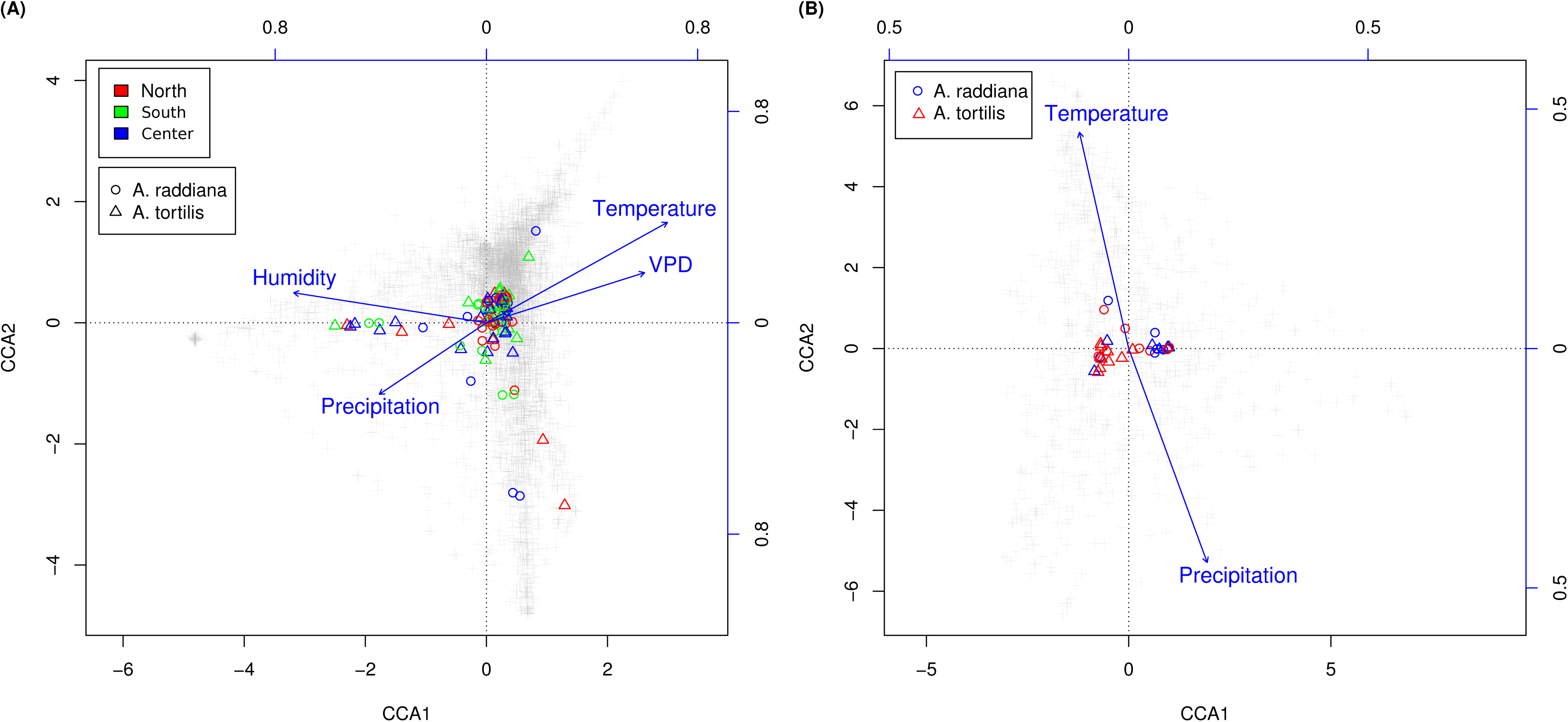
CCA ordination illustrating (A) exosphere bacterial community at north (red), south (green) and center (blue) canopy sides and (B) endophytic bacterial communities, for *A. raddiana* (circles) and *A. tortilis* (triangles) with significant abiotic factors affecting the bacterial communities.

To test for the major changes in bacterial species, the abundant OTU’s and their relative bacterial families were plotted as a heatmap for endophytic and exosphere south canopy side (Fig. 8). Results show that only few bacterial OTU’s were differentially abundant comparing exo and endophytic, or when comparing within the endophytic between *A. raddiana* and *A. tortilis*. The major differences occurred in 5 unclassified OTU’s belonging to *Bacillus*, *Comamonadaceae*, *Geodematophilaceae* and *Micrococcaceae* bacterial families.

**Figure 8.**
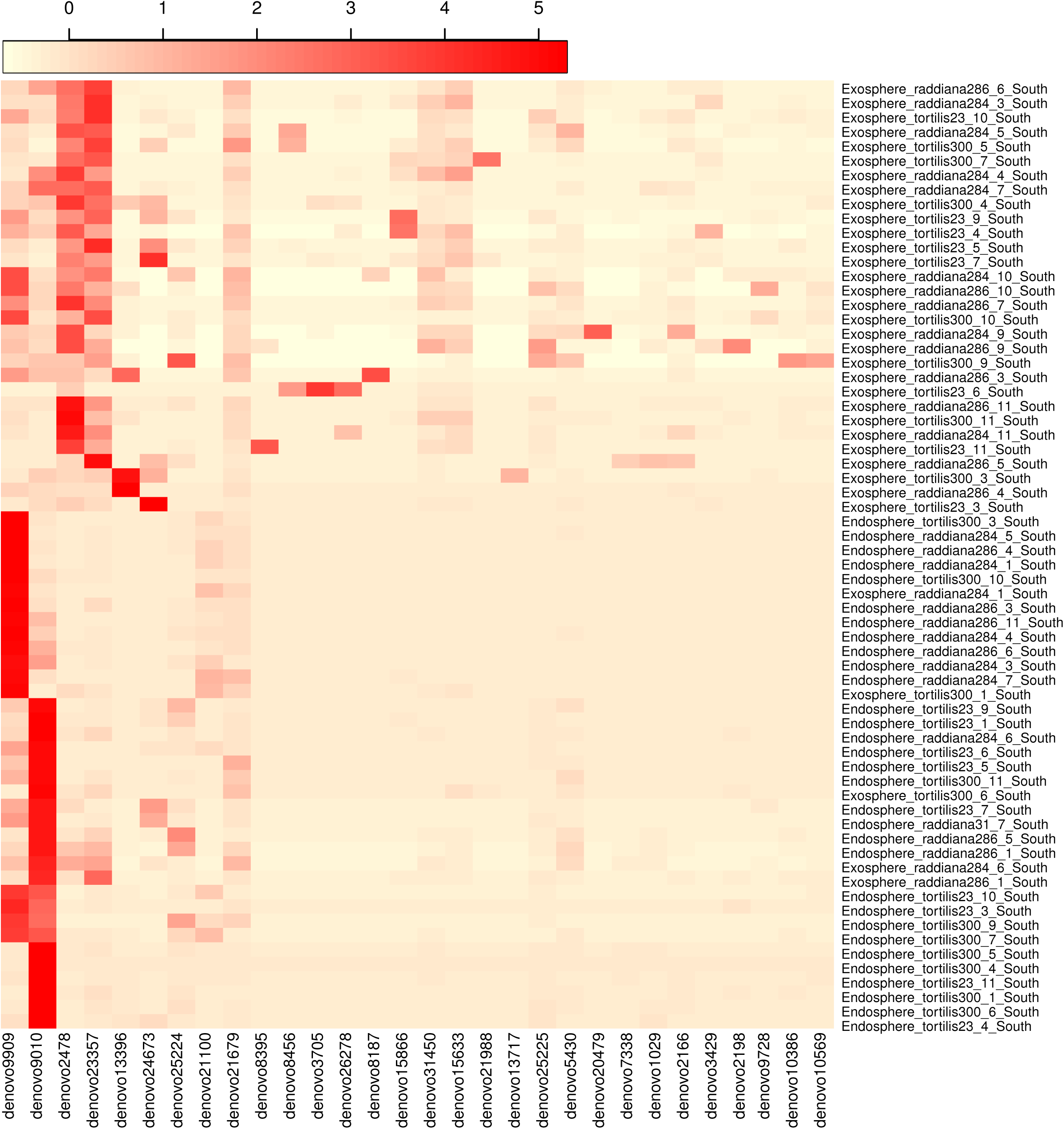
Heatmap showing the abundance of OTUs > 5% of the total bacterial communities (x-axis) for each of the sampled epiphytic and endophytic bacterial communities at south canopy side at different sampling months (1–12) during 2015.

## 3. Discussion

Aiming to improve our understanding of the functions that phyllosphere microbial communities might play in plants growing in extreme arid environments, we applied a high-resolution sampling scheme to studying the phyllosphere microbial communities of two desert keystone trees (*Acacia raddiana* and *Acacia tortilis*). We investigated both the endophytic and epiphytic bacterial communities to understand the: (i) intra- and inter-individual spatial variation of the microbial community within a tree (the variation within the same tree caused by different sides of the canopy, and the variation between neighboring trees of the same species sampled at the same time and site) (ii) host species variation (variation of the microbial community caused by the host (tree) species (i.e., *Acacia raddiana* compared with neighboring *Acacia tortilis*), (iii) temporal variation of the microbial community within the same tree species and canopy side (samples collected from the same trees but at different seasons).

Our results demonstrate that the epiphytic bacterial communities were more sensitive to changes in the external environmental conditions, compared with the endophytic bacterial communities that were more stable between different environmental conditions (e.g., seasons) but varied among host tree species. Surprisingly, up to 60% of the total bacterial communities (the combined endophytic and epiphytic microbiome populations) were unclassified below family level, highlighting the uniqueness of the microbiome associated with acacia trees in the arid environment in the Arava.

The epiphytic bacterial diversity was found to be significantly higher than the endophytic bacterial community (Table 2). In terms of the overall observed number of OTU’s, the epiphytic bacterial community was shown to have double the diversity compared to its endophytic bacterial community counterpart. Similar findings at early and late leaves development in *Origanum vulgare* also found the total number of colony-forming units (CFU) of endophytic communities (1.8 ± 0.1) was less than half of the CFU of epiphytic bacterial communities (5.0 ± 0.2) (34). However, our results contradict previous work on microbiomes associated with *Arabidopsis thaliana*) that showed epiphytic bacterial diversity indexes were lower than those measured for the associated endophytic bacterial communities (35). A recent study on the epiphytic and endophytic fungal diversity in leaves of olive trees growing in Mediterranean environments, showed that the epiphytic fungal communities had higher diversity indices compared to the endophytic diversity estimates (23). The fact that our epiphytic OTU diversity was higher than the endophytic diversity is particularly surprising, considering previous publication indicated that as the conditions inside the plant might be more favorable compared to the hostile conditions outside (36). This might explain the different findings comparing epiphytic and endophytic bacterial abundance and diversity in other studies, but in our case, both *A. raddiana* and *A. tortilis* had a lower epiphytic bacterial diversity compared to epiphytic bacterial diversity throughout the sampling month, including the hot and harsh conditions of the desert summer (Table S5). This discrepancy finding in our results and previously document findings could be unique to desert plants. Plants grown in desert environments are subjected to continuous stress conditions including increased salt concentration in endophytic compartments (37), decreasing stomatal conductance and increased concentration of abscisic acid (38) and many other metabolites and enzymes (39). These plant responses were shown to affect plants-microbiome colonization (34, 40, 41). Moreover, our results showed that the endophytic and epiphytic bacterial communities were significantly different from each other (Fig. 2A). In fact, endophytic but not epiphytic bacteria communities, differed between the two acacia species (Fig. 3A, 3B, 4). This potentially indicates that endophytic bacteria were horizontally transmitted and that they might be more affected by genotypic factors rather than abiotic factors (4, 5, 22).

Similar to other findings indicating the changes in bacterial communities in phyllosphere following different environmental and biotic factors (40, 41), our results show seasonality to be the major driver of community composition in epiphytic bacteria (Fig. 4A and Fig. 6), including a specific abiotic parameters such as; humidity, temperature, precipitation and VPD (Fig. 7A and B). While these results highlight the significant effect of temperature on both epiphytic and endophytic bacterial communities, the effect of microclimate (different canopy sides) on the epiphytic bacterial diversity (Table 3) and community composition (Fig. 5 and 6) showed no significant variation for the different canopy sides for both species. This could be explained by the difference between monthly temperature, humidity and precipitation hinders back these effects of canopy side variation.

We also showed that the bacterial community compositions found in this study, differ from other epiphytic or endophytic microbiome found in tropical, subtropical and temperate regions, which are mostly dominated by high abundance of *Alphaproteobacteria*, *Bacteroidetes* and *Acidobacteria* (1, 42, 43). In our study, the major differences between epiphytic and endophytic bacterial communities were due to the differential abundance of four major unclassified OTU’s belonging the bacterial families of *Bacillaceae* (Firmicutes phylum) and *Comamonadaceae* (Betaproteobacteria phylum) for the endosphere of *A. raddiana* and *A. tortilis*, respectively (Fig. 8). Other unclassified OTU’s belonging to the bacterial families of *Geodematophilaceae* and *Micrococcaceae* (both belonging to Actinobacteria phylum) were found in the exosphere bacterial communities (Fig. 8). These bacterial families were also found in other studies investigating extreme conditions that investigated the metagenomic signatures of *Tamarix* phyllosphere (8, 11, 44) and other desert shrubs (7), highlighting the importance and the relationship of these found bacterial communities in desert plants adaptation to arid environment (7). However, the exact link between these different bacterial group and their functional diversity is still to be investigated, such studies could shed the light of specific metabolites and enzymes that these adaptive bacterial group exhibit in such environment and at different stress conditions. Learning from the long co-evolved plants-microbiome form naturally occurring plant in harsh conditions is in vital under the current rate of climate change and the urgent need for new innovative solutions that can be learned from these interactions for more adaptive arid land agriculture.

## 4. Conclusion

The evolutionary relationship and interaction between plants and their microbiome is of high importance to their adaptation to extreme conditions. Changes in plants microbiome can affect plant development, growth and health. In this study we explored the relationship between naturally occurring desert plant and their microbiome along seasonal and abiotic conditions. These results shed light on the unique desert phyllosphere microbiome in mitigating stress conditions highlighting the importance of epiphytic and endophytic microbial communities which are driven by different genotypic and abiotic factors. Nevertheless, more studies utilizing the functional diversity of these different plants-microbiome interactions in arid climate is in vital with desertification and global warming processes in mind. The potential Agritech of these unique microbial communities calls for more research on this topic in the future exploring the functional diversity of each endo and epiphytic microbial communities alongside with plants metabolites at different stress conditions.

## 5. Materials and Methods

### 5.1. Study area and sampling scheme

This study was conducted in the Arava valley, a hyper-arid region along the Syrian-African rift in southern Israel and Jordan. The elevation of the area ranges from 230 m above sea level to 419 m below sea level (Fig. 1A). The climate in the Arava valley is hot and dry: 30-year average minimum, mean, and maximum air temperature of the hottest month was 26.2 °C, 33.2 °C, and 40.2 °C, respectively; average minimum, mean, and maximum air temperature of the coolest month were recorded as 9.1 °C, 14.4 °C, and 19.6 °C, respectively, and annual precipitation of only 20–70 mm is restricted to the period between October and May (32) with large year-to-year variations (45). The combination of the very high air temperatures and the very low relative humidity values of 6% can cause summer midday vapor pressure deficit (VPD) to reach up to 9 kPa (32). Vegetation in the region is usually confined to within wadis (ephemeral river beds (46)), where the main water supply comes from underground aquifers (47, 48) and winter flash floods (49). Multiple individual trees of *A. raddiana* and *A. tortilis* are scattered throughout the Sheizaf wadi (Fig. 1A), but never forming a continuous canopy. To investigate the effect of different canopy sides on phyllosphere microbiome, leaf samples were also collected from three different canopy sides (north, center and south; Fig. 1D).

Wadi Sheizaf is a dry sandy streambed at the northern edge of the Arava Valley, Israel (Fig. 1A; 30°44’N, 35°14’E; elevation −137 m). Meteorological data (air temperature and humidity logged every 3 hours) for this site were obtained from the Israeli Meteorological Service (IMS) for station 340528 at Hatzeva, located 7 km north of Wadi Sheizaf (Fig. 1C).

For sampling bacteria from acacia trees, two neighboring trees (>20 m away from each other) of *A. tortilis* (T023 and T300) and two neighboring trees of *A. raddiana* (R284 and R286) in Wadi Sheizaf, were sampled monthly between January and December 2015 for their North, South and Central canopy sides (Fig. 1D and Table S1). This sampling scheme was chosen to enable us to investigate the effect of having two different host (tree) species, in addition to the variation caused by the sampling season and the micro-climate effect (different canopy sites (central, north, and south-facing sides of tree) on the phyllosphere microbiome.

During all sampling campaigns, samples were collected from trees using sterile gloves (changed between each sample). Leaves (20-25 g fresh weight) were collected monthly (see Table S1 for exact dates) and inserted into 15 ml sterile tubes placed on ice. Upon reaching the laboratory (within <2 hrs) samples were moved to freezers (−20 °C) where they were kept until subjected to DNA extraction.

### 5.2. DNA extraction

All DNA extractions were performed using the MoBio 96 well plate PowerSoil DNA Isolation Kits (MO BIO Laboratories, California, USA). For exosphere “outside of plants leaves”, 0.15 g (FW) of leaves were weighed and placed in 1.5 ml Eppendorf tubes filled with 500 µl MoBio Power bead Solution and sonicated (DG-1300 Ultrasonic cleaner, MRC LAB, Israel) for 5 min and then the solution was transferred to the Power bead Tubes and the rest of the steps for DNA extraction were carried out following the manufacturer protocol. For the extraction of endophytic (“inside plant leaves”) microbial communities, leaves were washed using 1 ml of DNA/RNA free water three times to get rid of as much of the exosphere microbiome fraction. The washed leaves were then cut into small pieces using a sterile scalpel and placed into the MoBio 96 well Power bead plate for DNA extraction following the manufacturer’s protocol. All steps of DNA extraction were carried out in a sterile UV-hood (DNA/RNA UV-cleaner box, UVT-S-AR bioSan, Ornat, Israel) to reduce external contaminations. In every DNA extraction 96 well plate, DNA extraction negative controls were added by placing 200 µl of RNase free water (Sigma Aldrich, Israel). All samples were placed randomly in the DNA extraction plate to exclude any bias.

### 5.3. PCR, library preparation and Illumina sequencing

In order to obtain a better phylogenetic resolution and diversity estimate, a multiplex PCR using five different sets of the 16S rDNA genes was applied to cover about 1000bp of the 16S rRNA gene (Table S2).

First PCR (PCR-I) reactions were performed in triplicates, where each PCR-I reaction (total 25 µl) contained; 12.5 µl of KAPA HiFi HotStart ReadyMix (biosystems, Israel); 0.4 µl of equal v/v mixed primers forward and reverse primers (Table S2); 10 µl of molecular graded DDW (Sigma, Israel) and; 2 µl of (2-100 ng/µl) DNA template. PCR-I reactions were performed in Biometra thermal cycler (Biometra, TGradient 48) as the following: initial denaturation 95 °C for 2 min, followed by 35 cycles of 98 °C for 10 sec, 61 °C for 15 sec, and 72 °C for 7 sec. Ending the PCR-I routine was a final extension for 72 °C for 1 min. Upon completion of PCR-I, an electrophoresis gel was run to verify all the samples worked successfully. Following successful and verified amplification, triplicate samples were pooled together and were cleaned using Agencourt® AMPure XP (Beckman Coulter, Inc, Indianapolis, USA) bead solution based on manufacturer’s protocol.

#### 5.3.1. Library Preparation and Illumina sequencing

Library preparation was performed using a second PCR (PCR-II) to connect the Illumina linker, adapter and unique 8 base pair barcode for each sample (50). The PCR-II reactions were prepared by mixing 21 µl of KAPA HiFi HotStart ReadyMix (biosystems, Israel), 2 µl of mixed primers with Illumina adapter (Table S3), 12.6 µl of RNase free water (Sigma, Israel), 4 µl of each sample from the first PCR product with 2 µl of barcoded reverse primer, and placed in Biometra thermal cycler (Biometra, TGradient 48) as the following: initial denaturation 98 °C for 2 min, and then 8 cycles of 98 °C for 10 sec; 64 °C for 15 sec; 72 °C for 25 sec; and final extension of 72 °C for 5 min. Then all PCR-II product were pooled together and subjected to cleaning using Agencourt® AMPure XP (Beckman Coulter, Inc, Indianapolis, USA) bead solution based on manufacturer’s protocol, where 50 µl of pooled PCR-II product were cleaned using 1:1 ratio with the bead solution for more conservative size exclusion of fragments less than 200 bp, and at the final step, 50μl of DDW with 10 mM Tris [pH = 8.5] were added to each sample. This was followed by aliquoting 48 µl of the supernatant in to sterile PCR tubes and stored in −80 °C, while an additional 15 µl of the final product were sent to the Hebrew University (Jerusalem, Israel) where they were sequenced on full lane of 150PE Illumina Miseq platform.

### 5.4. Sequence analysis and quality control

A series of sequence quality control steps were applied before data analysis. These included the following steps: all samples were filtered for PhiX contamination using Bowtie2 (51); incomplete and low-quality sequences were removed by pairing the two reads using PEAR software (52); looking for ambiguous bases and miss merged sequences carried out using the MOTHUR Software V.1.36.1 (53). Following quality control, QIIME-1 software (54) was used.

Sequences were aligned, checked for chimeric sequences and clustered to different OTU’s (operational taxonomic unit) based on 97% sequence similarity, then the sequences were classified based on Greengenes database V13.8 (55), and an OTU table was generated. All sequences classified as f mitochondria, c Chloroplast, k Archaea and K Unclassified, were removed from the OTU table.

#### 5.4.1. OTU richness and diversity estimates

For each sample, four diversity estimates were measured; (i) observed number of OTU’s, (ii) Chao1 species’ diversity estimate (56), (iii) Shannon bacterial communities’ diversity (57), and (iv) microbial communities’ phylogenetic diversity (58). All these diversity metrics were calculated in QIIME-1 software (59) using the parallel_alpha_diversity.py command on the rarefaction subsamples to 10,000 sequences using multiple_rarefactions.py command.

#### 5.4.2. Assessment of community composition

From the obtained QIIME classified OTU table, each taxonomic group was allocated down to the genus level using summarize_taxa.py command in Qiime and relative abundance was set as the number of sequences affiliated with that taxonomic level divided by the total number of sequences. Relative abundances were plotted using R statistical software (60) where each phylum was assigned a distinguished color and all genera under the same phylum, were assigned to different shades of the same color.

#### 5.4.3. Statistical Analysis

Using R statistical software (60) all samples were analyzed based on the previously generated OTU table. Using VEGAN package (61) in R, non-parametric multidimensional scaling (NMDS) were used to produce ordination based on Bray-Curtis distance matrix using a total sum transformed matrix for the row OTU table (62, 63). Statistical comparisons were done based on analysis of the similarity between the matrices of different OTU’s table using ANOSIM command in R.

#### 5.4.4. Data availability

All curation steps, quality control and sequence analysis code uploaded to GitHub repository, including the metadata files and made publicly available (https://github.com/ashrafashhab/Desert-plant-microbiome). In addition, all curated sequences were joined into single fasta file and submitted to MG-RAST under a project link (https://www.mg-rast.org/linkin.cgi?project=mgp92155).

## 6. Acknowledgments

We thank Jewish Charitable Association in Israel (ICA) the funding agency for it generous support of this research that was awarded under grant number: 03-16-06A. We also thank Dr. Noam Shental from the Department of Computer Science at the Open University of Israel, for his generous support in preliminary data analysis, selection of 16s Primes and his insight in the experimental design. Also special thanks to Miss. Tal Galker from Arava Studio for developing Fig. 1 in the MS.

